# Imaginary scenes are represented in cortical alpha activity

**DOI:** 10.1101/2023.10.23.563249

**Authors:** Rico Stecher, Daniel Kaiser

## Abstract

Imagining natural scenes enables us to engage with a myriad of simulated environments. How do our brains generate such complex mental images? Recent research suggests that cortical alpha activity carries information about individual objects during visual imagery. However, it remains unclear if more complex imagined contents such as natural scenes are similarly represented in alpha activity. Here, we answer this question by decoding the contents of imagined scenes from rhythmic cortical activity patterns. In an EEG experiment, participants imagined natural scenes based on detailed written descriptions, which conveyed four complementary scene properties: openness, naturalness, clutter level and brightness. By conducting classification analyses on EEG power patterns across neural frequencies, we were able to decode both individual imagined scenes as well as their properties from the alpha band, showing that also the contents of complex visual images are represented in alpha rhythms. An additional cross-classification analysis between alpha power patterns during the imagery task and during a perception task, in which participants were presented images of the described scenes, showed that scene representations in the alpha band are shared between imagery and late stages of perception. This suggests that alpha activity mediates the top-down re-activation of scene-related visual contents during imagery.

## Introduction

Our ability to evoke mental images of natural scenes enriches our lives by giving shape to the worlds in our favorite novels or by enabling us to navigate the environment. Visual imagery is thought of as a top-down recall of sensory-memory information initiated in frontal cortex that reactivates cortical areas that are typically involved in visual perception (Pearson, 2019). For example, when imagining natural scenes, the scene network, a network of cortical areas that is typically active during scene perception, is re-engaged (Boccia et al., 2017; Johnson & Johnson, 2014; O’Craven & Kanwisher, 2000). Imagery recruits shared representations with perception across the entire visual hierarchy, although more extensively in high-level visual areas (Dijkstra et al., 2019; Pearson, 2019; Pearson et al., 2015). What neural mechanisms underlie this top-down re-instantiation of visual contents?

One possibility is that imagery-related information is encoded in neural rhythms that are involved in top-down processing. Previous research has provided evidence that alpha and beta rhythms carry top-down information in the visual cortex (Bastos et al., 2015; Chen et al., 2023; Fries, 2015; van Kerkoerle et al., 2014), making them likely candidates to play a role in imagery processing. In the past decades, the relation between alpha activity and visual imagery has been studied to quite an extent. While many studies have found a correlation between changes in alpha rhythms and visual imagery (e.g. Bartsch et al., 2015; Michel et al., 1994; Salenius et al., 1995; Short, 1953; Slatter, 1960), what exact information is encoded in these alpha rhythms has remained elusive for a long time. However, more recently popularized multivariate pattern analysis (MVPA; Grootswagers et al., 2017; Haynes, 2015) techniques have since enabled us to probe the representational content of alpha activity during visual imagery.

A recent EEG MVPA study (Xie et al., 2020) found that alpha oscillations contained representations of the visual contents of imagined objects. Interestingly, these alpha band representations were also shared with late time windows during perception. While this does provide evidence that alpha oscillations carry out the top-down re-instantiation of perceptual contents, the imagery task employed in this study (imagining isolated objects) is not very representative of the imagery tasks we typically perform in our daily lives. A lot of our everyday tasks like reading, spatial navigation or mental simulation of future events require us to imagine not just single objects, but complex natural scenes (Epstein, 2008; Mak et al., 2020; Schacter et al., 2017). Compared to isolated objects, perceptual processing of natural scenes requires several scene-specific steps such as the analysis of scene-diagnostic low-level image statistics (Groen et al., 2017), (global) scene properties (Park et al., 2015) and object arrangements (Kaiser et al., 2019; Võ, 2021). It is thus critical to understand if the top-down re-instantiation of scene information during imagery is also mediated by cortical alpha activity.

In the present study, we thus aimed to answer the question if natural scenes and their properties are represented in cortical alpha activity during visual imagery and if these representations are shared with perceptual processing. To that end, we conducted an EEG experiment in which participants imagined natural scenes based on a written description and viewed images of the same scenes in a separate task. The scenes varied in four properties which have previously been investigated in the scene literature: openness, naturalness, clutter level and brightness (Cichy et al., 2017; Harel et al., 2016). We employed frequency-resolved multivariate pattern classification to track the representations of scenes across neural rhythms, and imagery-perception cross-classification to investigate whether scene representations are similarly coded in rhythmic cortical activity during imagery and perception.

We show that imagined natural scenes and their properties are represented in cortical alpha activity and that scene representations in the alpha frequency band are partly shared between scene imagery and late stages of scene perception. Our results indicate that cortical alpha activity mediates the top-down re-instantiation of complex natural scenes during visual imagery.

## Results

In order to investigate the neural representations of imagined and perceived scenes and their properties in neural rhythms, we conducted two experimental tasks while participants’ neural activity was recorded via EEG (see Fig. 1c for a schematic of the tasks). In the imagery task, participants imagined natural scenes for 4000 ms based on a detailed, three-sentence description of each scene (see Fig. 1a for an example). The 16 described scenes varied independently in four properties: openness, naturalness, clutter level and brightness. After completing the imagery task, participants were asked to rate the imagined scenes in regard to the four properties on a scale from 1 to 7. These ratings confirmed that, at least on average, the properties of the imagined scenes aligned with those conveyed by the descriptions (see Fig. 1b). In the subsequent perception task, participants viewed images that matched the scene descriptions (3 images per scene; see Fig. 1a) with each image being presented for 1000 ms.

**Figure. 1.**
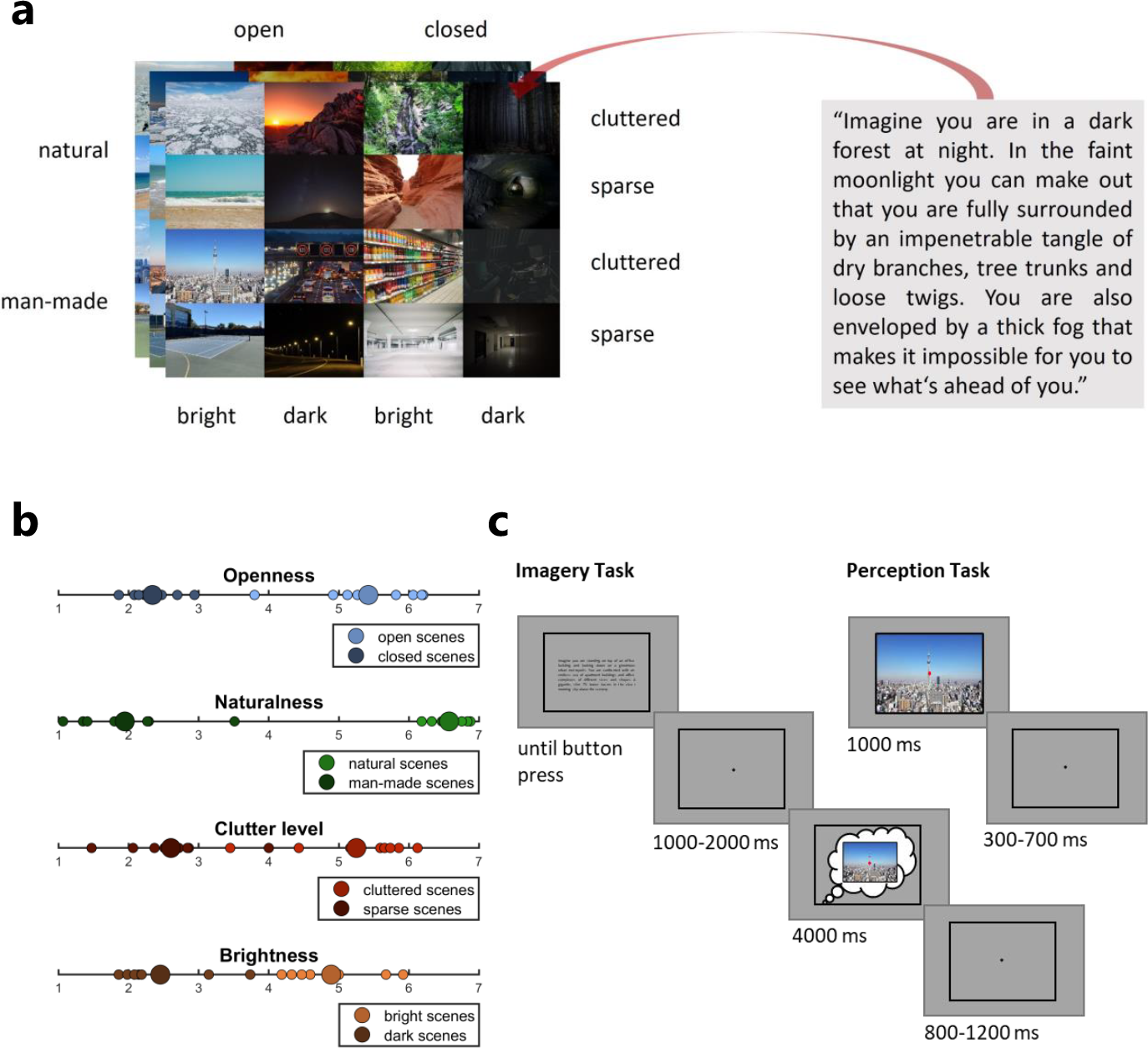
Stimuli and Paradigm. a) Stimuli used in the EEG experiment. Participants imagined 16 natural scenes according to detailed three-sentence descriptions (right) and in a separate task viewed images that matched these descriptions (three per scene; left). The scenes varied in their openness, naturalness, clutter level and brightness. b) Average ratings of all four properties for each imagined scene. After finishing the imagery task, participants rated each imagined scene on a scale from 1 to 7 regarding the four investigated properties openness, naturalness, clutter level and brightness. On average, property ratings of imagined scenes aligned with the properties conveyed by the scene descriptions (e.g. the mental images of open scenes were also rated as high on the openness dimension and those of closed scenes were rated as low). Large circles: property category mean rating. Small circles: mean rating of individual scenes in the respective property category. c) Experimental tasks. In the imagery task, participants were presented with a scene description surrounded by a black frame until they proceeded with a button press. A black fixation dot appeared and after a jittered interval of 1000-2000 ms, the fixation dot turned red which was their cue to imagine the scene within the surrounding frame. They were instructed to maintain the mental image of the scene while fixating the red dot until it turned black again after 4000 ms. In the perception task, participants were presented images for 1000 ms that matched the scene descriptions in the imagery task and were only tasked with attentively viewing them.

### Mean pairwise scene decoding

To identify at which neural frequencies information related to individual imagined scenes can be found, we conducted a mean pairwise frequency searchlight decoding analysis. We transformed the EEG signals across the entire imagery period into the frequency domain and trained classifiers to distinguish between each possible pair of imagined scenes based on power patterns across channels at each frequency from 4-30 Hz (see Fig. 2a for a schematic). Averaging the pairwise decoding accuracies across pairs yielded a measure of information content regarding the individual imagined scenes at each frequency. We found that the individual imagined scenes could be discriminated the best in the alpha frequency range (see Fig. 3a), with significant mean pairwise scene decoding from 8-13 Hz, peaking at 11 Hz (p<0.001). There was also some weaker, but sporadically significant mean pairwise scene decoding in the beta band (significant at 18 and 21 Hz), indicating that some information might also be contained there. These results suggest that individual imagined scenes are represented most prominently in the alpha frequency band.

**Figure. 2.**
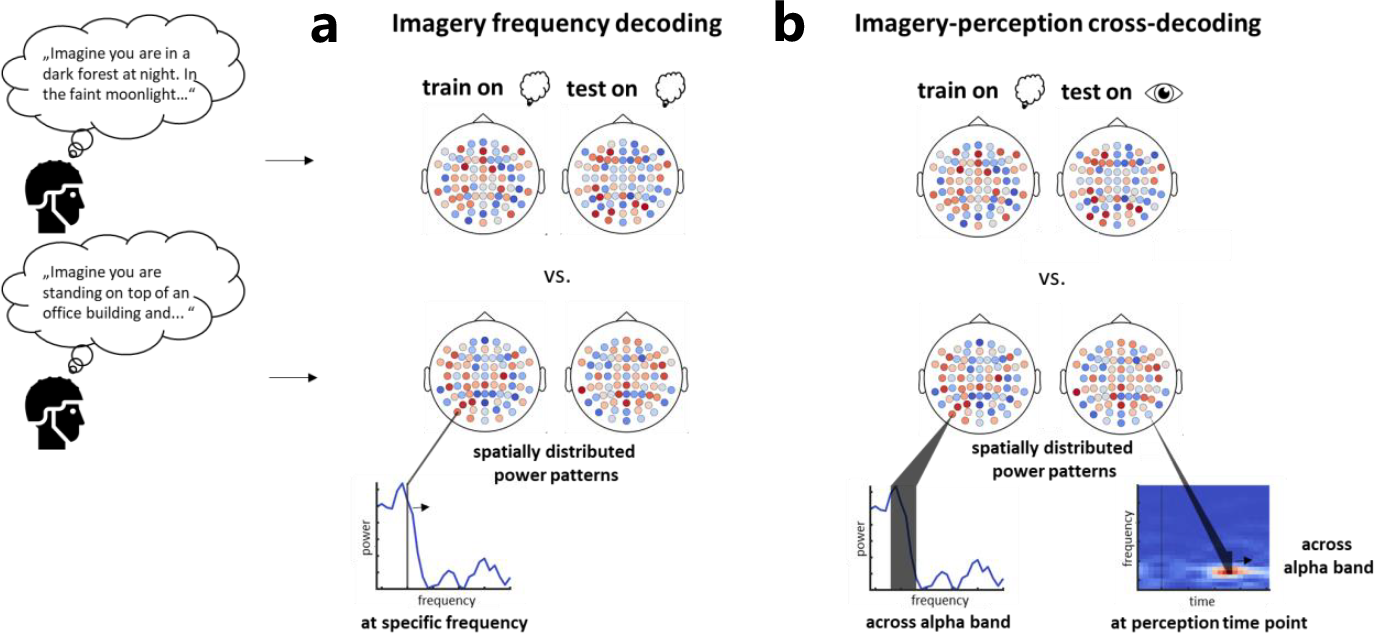
Decoding approaches. We conducted two main decoding analyses. In the imagery frequency decoding analysis (a), we investigated rhythmic scene representations during imagery by training classifiers on power patterns across all channels at each frequency from 4-30 Hz in the frequency-resolved imagery EEG data. In the imagery-perception cross-decoding analysis (b), we investigated shared scene representations between imagery and perception in the alpha band, by training the classifiers on alpha power patterns across all channels in the frequency-resolved imagery data and testing them on alpha power patterns across all channels at each time point in the time-frequency-resolved perception data and vice versa. We employed two decoding schemes in both analyses: a mean pairwise scene decoding and a scene property decoding. In the mean pairwise scene decoding, classifiers were trained to distinguish between each possible pair of scenes and decoding accuracies were averaged across each scene pair, as a measure of neural discriminability among individual scenes. In the scene property decoding, classifiers were trained to predict for each property in which of two property categories a given scene belonged. In an additional control analysis, we re-conducted the scene property (cross-) decoding analyses by training classifiers to discriminate between mock property categories with randomly assigned scenes as a measure of neural discriminability between property categories based on individual scene features.

**Figure. 3.**
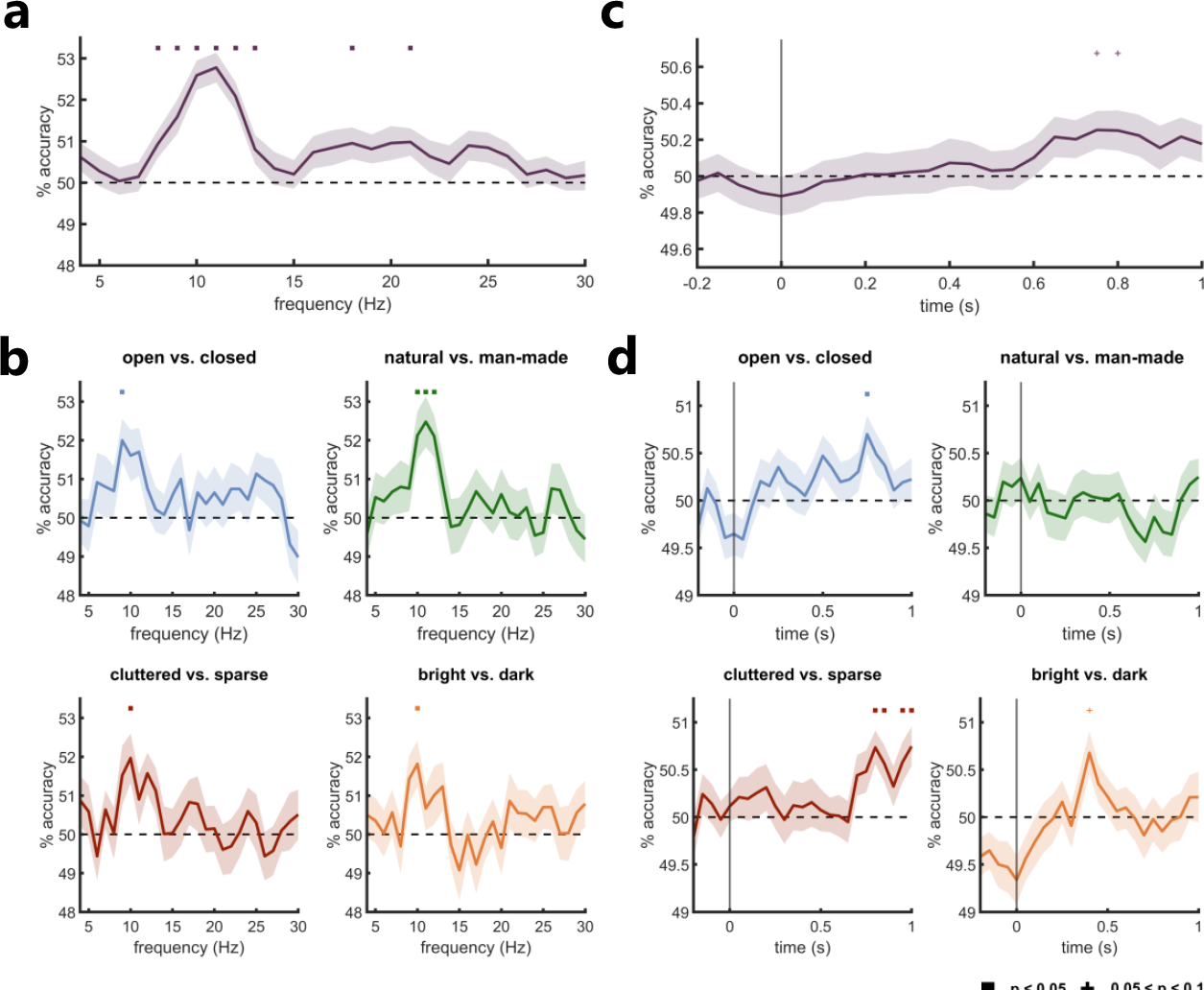
Decoding of imagined scenes and their properties based on spatially distributed power patterns in rhythmic neural activity. a) Mean pairwise decoding of imagined scenes at each frequency from 4-30 Hz. Individual imagined scenes could be discriminated the best from alpha band activity (8-13 Hz). b) Scene property decoding. All properties could exclusively be decoded from the alpha frequency band. c) Mean pairwise imagery-perception scene cross-decoding in the alpha frequency band at each time point during scene perception. A marginally significant trend suggests that there are shared representations of individual scenes in the alpha frequency band between imagery and late stages of perception. d) Imagery-perception scene property cross-decoding in the alpha frequency band at each time point during scene perception. For openness, clutter level and brightness (brightness being only marginally significant) we found evidence of shared representations in the alpha frequency band between imagery and late stages of perception. Error margins reflect the standard error of the mean. Square markers indicate significance at p<0.05, cross markers indicate marginal significance at p<0.1 (both corrected for multiple comparisons).

### Scene property decoding

We further examined how the four investigated properties (openness, naturalness, clutter level and brightness) of the imagined scenes are represented across the frequency domain. Using the same frequency searchlight approach as above, we had classifiers predict for each property which property category the imagined scene belonged to (e.g., for naturalness if the scene was natural or man-made). This analysis revealed that all four properties could exclusively be decoded from the alpha band (see Fig. 3b). The decoding accuracy for each property peaked at around 10 Hz (openness: 9 Hz, p=0.009; naturalness: 11 Hz; p<0.001, clutter level: 10 Hz, p=0.038; brightness: 10 Hz, p=0.031). Aligning with our decoding analysis of the individual scenes, these findings suggest that the properties of imagined scenes are also represented in cortical alpha activity.

### Imagery-perception cross-decoding in the alpha frequency band

Next, we investigated if any of the of representations of the individual scenes or scene properties we found in the alpha band are shared between imagery and different stages of the perceptual processing hierarchy. To that end, we conducted the same mean pairwise and property decoding analyses, but trained the classifiers on alpha power patterns in the frequency-resolved imagery data and tested them on alpha power patterns at each time point in the time-frequency-resolved perception data and vice versa (see Fig. 2b for a schematic). We chose to assess these shared alpha representations across the entire imagery period while maintaining temporal resolution for the perceptual data in order to increase the power of the analysis, since imagery representations were found to be relatively invariable across time (Corriveau et al., 2023; Dijkstra et al., 2018; Xie et al., 2020), whereas perceptual representations are thought to be more temporally variable (Carlson et al., 2011; Dijkstra et al., 2018; Singer et al., 2023). This yielded a time-resolved measure of shared representations in the alpha band between scene imagery and each stage in the processing hierarchy of scene perception, in which temporal representational variations index the procession of perceptual processing (King & Dehaene, 2014). In the mean pairwise scene cross-decoding analysis, we identified an increase in mean pairwise scene cross-decoding accuracy starting at around 600 ms during perceptual processing that was marginally significant (p=0.068 at peak) at 750-800 ms (see Fig. 3c). This trend suggests that there are representations of individual scenes in the alpha band that are shared between scene imagery and late stages of scene perception. In the property cross-decoding analysis, we found relatively low but significant cross-classification performance for openness at 750 ms (p=0.012) and for clutter level at 800-850 ms (p=0.002 at peak) as well as 950-1000 ms (p=0.008 at peak) in the perceptual processing hierarchy (see Fig. 3d). There also was marginally significant (p=0.057) cross-decoding performance for brightness at 400 ms, but no significant cross-decoding performance for naturalness. The peak in cross-decoding accuracy for openness was very similar to the first peak in cross-decoding accuracy for clutter level and both aligned temporally almost perfectly. They both also temporally overlapped with the marginally significant peak in the mean pairwise cross-decoding peak for the individual scenes. The results of our property cross-decoding analysis indicate that scene imagery shares representations with late scene perception in the alpha band, at least for some properties. We exploratively conducted all imagery-perception cross-decoding analyses in the theta and beta bands as well, but there was no solid evidence of shared scene representations in those frequency bands (see supplementary Fig. S2).

### Shuffled property decoding

In a further control analysis, we assessed to what extent the representations the classifiers utilized during scene property decoding encode differences in individual scene features or property category information. We performed a shuffled property decoding analysis for both the imagery property decoding and imagery-perception property cross-decoding in which all scenes were randomly assigned to two mock property categories for all possible permutations and classifiers were trained to distinguish between these categories. Since property category information is randomized in this decoding scheme, the classifiers should be limited to differentiate between these mock categories based on the features of the individual scenes they happen to encompass. If the property decoding did not only use differences in individual scenes, but also more abstract information on property categories, shuffled property decoding performance should be reduced in comparison to the original property decoding performance, since the classifiers had no access to this additional source of information in the shuffled analysis. To test this, we compared the peak (cross-)decoding accuracy for each property that was discriminable in the original analyses to the shuffled property (cross-)decoding accuracy at the respective frequency or time point.

For the imagery property decoding analysis, we found higher peak decoding accuracies compared to the shuffled property decoding accuracies for all properties (see Fig. 4a). This difference was significant for openness (p=0.015) and naturalness (p=0.031), marginally significant for clutter level (p=0.072) and not significant for brightness (p=0.102). This implies that for openness, naturalness and potentially clutter level, there are property category representations in the alpha band during scene imagery. We also conducted the shuffled property decoding at each frequency during imagery (see supplementary Fig. S3). While slightly lower in overall accuracy, the decoding performance profile looks strikingly similar to that in our mean pairwise scene decoding, further corroborating that the shuffled property decoding mainly reflects neural discriminability based on individual scene information.

**Figure. 4.**
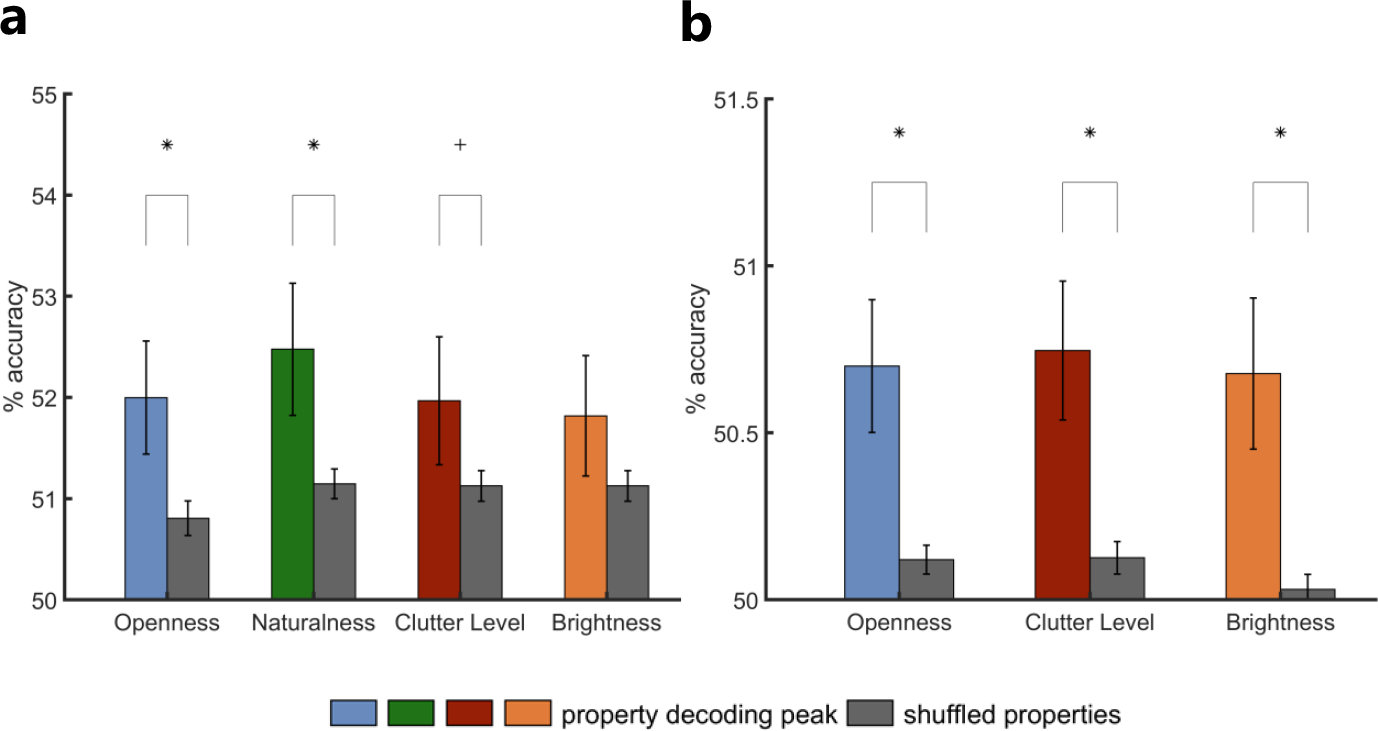
Comparison between peak scene property decoding accuracies and decoding accuracies with randomized property assignment (shuffled property decoding). a) Comparison at the peak decoding frequency for each property in the imagery property decoding. Property decoding accuracies exceeded shuffled decoding accuracies for all properties. This was significant for openness, naturalness and marginally for clutter level, suggesting that there are representations of property category information of these properties in the alpha band. b) Comparison at the peak imagery-perception property cross-decoding time points for all properties that were discriminable in the original analysis. For each property, property cross-decoding accuracies significantly exceeded the shuffled accuracies, suggesting that there are property category representations in the alpha band that are shared between scene imagery and late stages of scene perception. Error bars reflect the standard error of the mean. Asterisks indicate significance at p<0.05, cross markers indicate marginal significance at p<0.1.

For the imagery-perception property cross-decoding analysis, we conducted the comparison between property cross-decoding and shuffled property cross-decoding for all properties except naturalness, since we did not find any interpretable cross-decoding accuracy peak for this property. We found higher cross-decoding accuracies compared to the shuffled property cross-decoding accuracies for all three investigated properties (see Fig. 4b). This was significant for all properties (openness: p=0.008, clutter level: p=0.004, brightness: p=0.006), suggesting that there are category representations of these properties in the alpha band that are shared between imagery and late stages of perceptual processing. When conducting the shuffled property cross-decoding at each time point during perception (see supplementary Fig. S3), we found significant above-chance cross-decoding performance starting at about 750 ms, providing further evidence of shared scene representations between imagery and late perception in the alpha band.

## Discussion

In the present study, we investigated the representations of imagined and perceived natural scenes in rhythmic cortical activity. We found, as hypothesized, that both individual scenes as well as scene properties are represented in cortical alpha activity during visual imagery. We also found evidence that scene representations in the alpha frequency band are partly shared between imagery and late stages of perceptual processing.

These results indicate that the top-down reactivation of scene representations during visual imagery is enabled by cortical alpha activity. This aligns well with studies showing that alpha rhythms play a role in visual imagery (e.g. Bartsch et al., 2015; Michel et al., 1994), and specifically with the notion that imagery-related alpha oscillations are a top-down signal that represents the imagined visual contents (Xie et al., 2020). In more general terms, our results also support theories that postulate that top-down information flows are mediated by alpha dynamics in visual cortex (e.g. Fries, 2015).

Our decoding analyses revealed that all four investigated scene properties were discriminable from cortical alpha activity during visual imagery and that for all scene properties except naturalness (and brightness being only marginally significant) these alpha representations were shared with late stages of scene perception. Comparing the initial imagery property decoding to a decoding scheme in which property categories were randomized showed that there were genuine and abstract representations of scene properties in the alpha band, which were observed for all properties except brightness during imagery (clutter level being only marginally significant). Extending this comparison to the imagery-perception cross-decoding, we found evidence of shared alpha representations of abstract property information for openness, clutter level and brightness. These results suggest that, during imagery, alpha activity enables the top-down reactivation of scene property representations, some of which are shared with late stages of scene perception. This further implies that the representational division into (global) scene properties found during perception (Greene & Oliva, 2009) also holds during imagery. One exception was, however, that we did not find any shared alpha representations for naturalness. A feasible explanation might be that, while participants rated the naturalness of the imagined scenes as expected, the natural and man-made contents they imagined might have differed from the natural and man-made contents in the images they viewed (e.g. different types of natural or man-made objects), resulting in different neural representations being recruited during imagery and perception. The properties for which we did find evidence of shared representations (openness, clutter level and brightness) are much less dependent on the specific types of imagined objects.

We found evidence of shared scene representations in the alpha band between imagery and perception from around 400 ms (for brightness) until 1000 ms (for clutter level) after stimulus onset. This is in alignment with previous studies employing cross-decoding techniques which have also reported late shared representations during perception. Xie et al. (2020), who originally found shared alpha representations between object imagery and perception, also reported late timings during perceptual processing with the strongest correspondence with imagery for perception emerging after 400 ms. Dijkstra et al. (2018) reported shared representations between imagery and perception up until 1000 ms during perception. Why would imagery reactivate representations that occur so late during perceptual processing? One potential explanation is that imagery and perception share fewer representations in low-level and more in high-level visual areas (Dijkstra et al., 2019), making it more likely that shared representations occur during later perceptual processing. This can be explained by the prominent conceptualization of imagery as a reverse reactivation of the perceptual hierarchy starting from high-level visual cortex (Dijkstra et al., 2020; Linde-Domingo et al., 2019). Following this notion, the representational format of cortical brain areas in late stages of perceptual processing (i.e. high-level visual cortex) is thought to be more similar to those in imagery since they are closer to the trigger source of the imagery signal (Pearson, 2019). In alignment with this, Xie et al. found that the late shared alpha representations were best explained by complex visual features analyzed in high-level visual cortex. Thus, the shared alpha representations in our results might also reflect late processing in high-level visual areas. This is supported by our shuffled property control analysis that yielded that some of the shared property representations in the alpha band encode category information, which is typically represented in high-level visual cortex (Grill-Spector & Weiner, 2014).

However, even if late shared representations between imagery and perception are not unexpected, the timings of our results are still quite late, given that processing of scenes and (global) scene properties (specifically openness, naturalness and clutter level) has been shown to be rapid and already occurs within the first 250 ms after stimulus onset (Cichy et al., 2017; Groen et al., 2013; Hansen et al., 2018; Harel et al., 2016; Lowe et al., 2018). Since most research on the temporal dynamics of scene processing has focused on comparatively early neural signatures (Harel et al., 2022), what happens during such very late stages of scene processing is still largely unknown. Given that the perceived scene images in our study were presented throughout the entire analysis time window, a possible explanation is that the alpha representations scene perception shares with scene imagery in our data reflect recurrent processing of the scenes and their properties after the first feed-forward sweep (Dijkstra et al., 2020). During recurrent processing, the perceptual representational format might be altered in a way that makes it more similar to imagery representations. Future studies could clarify to what extent recurrent processes shape the late representations in perception that generalize to imagery.

A final caveat in our results are the low decoding accuracies. Imagery-related brain signals tend to have a low signal-to-noise-ratio (e.g. Shatek et al., 2019) which results in lower decoding accuracies in imagery studies that employ MVPA (e.g. Dijkstra et al., 2018; Xie et al., 2020). Furthermore, our imagery task was designed to ensure that the imagined scenes sufficiently differed from the perceived scenes in terms of their low-level features. We had participants imagine the scenes based on descriptions that allow for variability in the generated mental images and only presented them the actual images after, so that if we did find shared representations between imagery and perception, they would not be based on similarities in low-level features. However, a side effect of this might have been reduced cross-decoding performance since the classifiers could not exploit such low-level features to a great extent. In addition, due to the relatively long trial duration, we only had 192 imagery trials of training data per participant, which further limited classifier performance. Nevertheless, low decoding accuracies can still constitute meaningful effects, indicating that information is represented consistently in neural response patterns across participants (Hebart & Baker, 2018; Robinson et al., 2023).

Overall, our results suggest that the top-down reactivation of scene representations during visual imagery is mediated by cortical alpha activity and that the re-instantiated alpha representations are partly shared with late stages of scene perception. They show that alpha dynamics are not only critical for generating mental images of individual objects, but also mediate the creation of complex natural environments in our mind’s eye.

## Methods

### Participants

50 participants (25 male; mean age=25.74 years, SD=6.31) with normal or corrected-to-normal eyesight took part in the experiment. One participant was excluded from all analyses because they did not complete the imagery task due to a technical error during the EEG recording. During recruitment, participants filled in a German translation of the Vividness of Visual Imagery Questionnaire (VVIQ; Marks, 1973), a common measure of a person’s aptitude at evoking mental images, on Limesurvey (https://www.limesurvey.org/en/). The scale of the VVIQ was reversed so that higher scores indicate better imagery performance and participants were only allowed to take part if they had a VVIQ score of at least 24/80, since scoring lower would constitute moderate to severe aphantasia (Zeman et al., 2020). They provided written informed consent and received monetary compensation. The study was approved by the ethics committee of the Julius-Liebig-University Gießen und was in accordance with the 6^th^ Declaration of Helsinki.

### Stimuli

Participants imagined and viewed 16 naturalistic scenes which independently varied in four properties: openness (8 open and 8 closed scenes), naturalness (8 natural and 8 man-made scenes), clutter level (8 cluttered and 8 sparse scenes) and brightness (8 bright and 8 dark scenes) (see Fig. 1a). During the imagery task, participants visualized each scene based on a three-sentence description in German (see Fig. 1a for an example and supplementary Table S1 for a list of all descriptions with English translation). The descriptions were detailed (mean word count=50.94, SD=5.23) in order facilitate evoking rich mental images of the scenes as well as to ensure that the visualized scenes were as similar in properties to the described scenes (and thus the scenes used in the perception task, see below) as possible. In the perception task, participants viewed color images of scenes that matched the descriptions in the imagery task. Each participant was presented with three different images per scene description, in order to account for the variability in imagined scenes when assessing shared neural representations between imagery and perception later on. The total of 48 images (16 scenes x 3 images; see supplementary Fig. S1 for all images) was taken from Google Images, cropped and resized to a resolution of 800 x 600 pixels (21° horizontal visual angle). Brightness and contrast were adjusted where necessary to accentuate the desired lighting conditions (bright vs. dark) or enhance visibility.

### Experiment design and procedure

The experiment consisted of two tasks (see Fig. 1c). Participants first performed the imagery task, in which they had to imagine scenes according to three-sentence descriptions (see Fig. 1a) while their EEG was recorded. At the beginning of each trial, participants were presented with a scene description at the center of the screen enclosed by a black frame (21° horizontal visual angle). The frame was visible throughout the entire trial to avoid evoking neural responses related to its onset. Participants were instructed to attentively read the description and try to identify all important aspects of the scene. They had an unlimited amount of time, but were asked to only take as long as needed, especially once they were more familiar with each description after multiple trials. Once they had familiarized themselves with the description, they pressed the spacebar and a black fixation dot appeared at the center of the screen. After a randomly jittered interval of 1000-2000 ms, the fixation dot turned red. This served as the cue to imagine the scene within the black frame that surrounded the red fixation dot. Participants were instructed to maintain the mental image of the scene while fixating the red dot until it turned black again after 4000 ms. Finally, a randomly jittered inter-trial interval (ITI) between 800-1200 ms followed, throughout which the black frame and fixation dot remained visible on screen. Before performing the imagery task, participants completed 16 practice trials (containing each scene once) which they underwent while their EEG cap was prepared. During the subsequent experiment, they were asked to imagine each of the 16 scenes 12 times, resulting in 192 trials (96 per property category). Trials were separated into 12 blocks. In each block, participants imagined all scenes once in random order. After performing the imagery task, participants filled in a questionnaire on Limesurvey in which they had to rate the mental image they had of each of the 16 scenes in respect to the four scene properties (openness, naturalness, clutter level, brightness) on a Likert scale from 1 to 7. We explicitly told the participants to make these ratings purely based on their mental images, even if they differed from the scenes in the descriptions. These ratings suggest that, at least on average, the properties of the scenes imagined by the participants aligned with the properties the scene descriptions intended to convey (see Fig. 1b for a plot of the ratings). After finishing the questionnaire, the participants took part in the perception task. Here, they viewed images of scenes that matched the scene descriptions in the imagery task while their EEG was again recorded. In each trial, a single image was shown at the center of the screen for 1000 ms. To make both tasks visually as similar as possible, we presented each stimulus in the perception task within the same black frame as in the imagery task and there also was a red fixation dot visible at the center of the image which the participants were instructed to attend continuously. Between each stimulus presentation, there was a jittered ITI of 300-700 ms in which again the black fixation dot surrounded by the black frame was visible. Each of the 48 images (see Stimuli) was presented 20 times for a total of 960 trials (480 per property category), except for one participant who only completed 416 trials (after balancing across categories) due to a technical error during the EEG recording. The trial order was fully randomized. Both imagery and perception tasks were interspersed by numerous self-paced breaks. The entire experiment, including EEG preparation, took between 2.5 to 3.5 hours. Stimulus presentation was controlled using Psychtoolbox (Brainard, 1997).

### EEG data acquisition and preprocessing

EEG data was acquired using an Easycap system with 64 channels and a Brain Products amplifier. The data was recorded at a sample rate of 500 Hz with Fz as the reference. The electrode arrangement followed the standard 10-10 system. All preprocessing was conducted using FieldTrip (Oostenveld et al., 2011). EEG was high- and low-pass filtered between 1-90 Hz, band-stop filtered to remove 50 Hz line noise, epoched between -1000 ms and 5000 ms for the imagery data and -1000 ms and 2000 ms for the perception data and baseline-corrected with a baseline window of 500 ms for imagery and 200 ms for perception. The EEG signal was then downsampled to 200 Hz. Noisy channels were removed by calculating the variance of each channel and rejecting outlier channels on this metric through visual inspection. Finally, independent component analysis (ICA) was applied to the EEG data and eye artifact components were removed through visual inspection.

### Frequency decomposition

All of our frequency decompositions were conducted separately for each trial and each channel. We transformed the EEG signals within the entire 4000 ms imagery period into the frequency domain. For the perception data, the period from -200 to 1000 ms was transformed into the time-frequency domain using a fixed-size 500 ms sliding window with 50 ms steps. For both decompositions, we utilized multitapers (15 DPSS tapers for imagery and 3 for perception) with constant 2 Hz frequency smoothing as implemented in FieldTrip. We chose multitapers in particular to increase power in our frequency-based analyses. Imagery data tends to be noisy (e.g. Shatek et al., 2019; Xie et al., 2020) and the multitaper approach typically increases the signal-to-noise ratio in frequency-resolved data at the expense of increased temporal and frequency smoothing (Cohen, 2014). In addition, we decided to omit temporal resolution of the frequency decomposition of the imagery data while maintaining it for the perception data in order to further boost power, since imagery representations have been shown to be relatively invariable across time (Corriveau et al., 2023; Dijkstra et al., 2018; Xie et al., 2020) while perceptual representations have been shown to be temporally variable as a function of the different processing stages in the visual hierarchy (Carlson et al., 2011; Dijkstra et al., 2018; King & Dehaene, 2014; Singer et al., 2023). Extracted frequencies ranged from 4-30 Hz, thus covering the theta (4-7 Hz), alpha (8-13 Hz) and beta (14-30 Hz) frequency bands.

### Decoding analyses

All decoding analyses were conducted using CoSMoMVPA (Oosterhof et al., 2016). We employed linear discriminant analysis (LDA) classifiers which were trained within-subject on power patterns across all channels (see Fig. 2).

In order to assess how the individual imagined scenes and their properties are represented in neural activity patterns, we employed two decoding approaches and solely altered the features on which we conducted these analyses to answer different questions. The first approach was a mean pairwise scene decoding analysis in which classifiers were trained to discriminate between each possible pair of scenes and the resulting pairwise decoding accuracies were averaged across pairs, yielding a measure of discriminability among individual scenes from neural responses. The second approach was a scene property decoding analysis in which classifiers had to distinguish for each property in which of two property categories a scene belonged (e.g. for naturalness, if the imagined scene was a natural or a man-made scene).

First, we assessed at which neural frequencies scene information is represented by running the mean pairwise scene decoding and the scene property decoding on power patterns at each individual frequency from 4-30 Hz in the frequency-resolved imagery EEG data (see Fig. 2a). Classifiers were trained using a leave-one-trial-out cross-validation scheme in which one trial per stimulus was left out to avoid imbalance between conditions. Decoding accuracies were calculated as the mean of all cross-validation fold accuracies. For the mean pairwise scene decoding, this resulted in one mean pairwise scene decoding accuracy at each frequency for each participant. For the scene property decoding, this yielded one decoding accuracy at each frequency for each property and each participant.

Second, we examined if there are scene representations in the alpha frequency band that are shared between imagery and different stages of perceptual processing across time. We again conducted the mean pairwise and property decoding analyses, but trained the classifiers on power patterns across the entire alpha frequency range (8-13 Hz) in the frequency-resolved imagery data and tested them on the alpha power patterns at each time point in the time-frequency resolved perception data and vice versa (see on Fig. 2b). The decoding accuracies of both train-test directions were averaged, which resulted in one mean pairwise cross-decoding accuracy time course as well as four property category cross-decoding accuracy time courses for each participant. We also exploratively conducted all imagery-perception cross-decoding analyses in the theta and beta bands (see supplementary Fig. S2).

Finally, it is possible that during the property decoding analyses the classifiers did not utilize property category information, but just exploited differences in features of individual imagined and perceived scenes. To investigate this, we conducted a shuffled property decoding analysis in which we estimated how well the classifiers perform if they are constrained to individual scene feature information, without access to property category information and compared this performance to the original property decoding. Within each participant, the 16 scenes were randomly assigned to two mock property categories for all possible 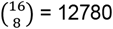 permutations and at each permutation, classifiers were trained to distinguish between the property categories. Decoding accuracies were then averaged across permutations. Since in this decoding scheme the property category information was randomized, classifiers were limited to discriminate based on differences in the individual scene features in each category. If the property decoding only exploited individual scene features, the shuffled property decoding performance should be identical or highly similar to it. If, however, property category information was also used for property discrimination in the original analysis, the shuffled property decoding performance should be reduced in comparison since the classifiers in the shuffled analysis had no access to this additional source of information. We applied the shuffled property decoding scheme to both the frequency-resolved imagery property decoding and our imagery-perception property cross-decoding. We tested the difference between property decoding and shuffled property decoding by assessing if the peak decoding accuracy of each property in the imagery property decoding and imagery-perception property cross-decoding is greater than the decoding accuracy at the respective frequency or time point in the shuffled property decoding. For the imagery-perception cross-decoding, this comparison was omitted for naturalness since we found no interpretable above-chance cross-decoding performance for this property in the original analysis. We also conducted the shuffled property decoding across all frequencies in the imagery property decoding and all time points in the imagery-perception property cross-decoding (see supplementary Fig. S3).

We investigated the temporal dynamics of imaginary scene representations as well. However, when conducting the aforementioned mean pairwise and property decoding analyses on broadband EEG responses at each time point, consistent with previous imagery studies (Shatek et al., 2019; Xie et al., 2020), we did not find robust above-chance decoding performance (see supplementary Fig. S4).

### Statistical testing

Decoding accuracies in all frequency-resolved and time-resolved analyses were tested against chance level (50%) using threshold-free cluster enhancement (TFCE; Smith & Nichols, 2009) as implemented in CoSMoMVPA. Multiple comparison correction was conducted by comparing actual TFCE statistics to a null distribution of maximum TFCE statistics, estimated using a permutation test with 10,000 sign permutations. The resulting z-scores were converted to p-values and thresholded at p<0.05 (one-tailed). In our time-resolved analyses, only the post-stimulus time points were tested for significance. We compared the peak property (cross-)decoding accuracies to the shuffled property (cross-)decoding accuracies at the respective frequency or time point using paired, one-tailed Wilcoxon signed rank tests. All statistical tests were conducted on the full sample of n=49.

## Data availability

Data and analysis code of the main analyses are openly available from our OSF repository: https://osf.io/vxhtw/.

## Additional information

The authors declare no conflicts of interest.

## Author contributions

R.S.: Conceptualization, Methodology, Data curation, Investigation, Formal analysis, Visualization, Writing — original draft, Project administration, Writing — review and editing

D.K.: Conceptualization, Methodology, Supervision, Project administration, Funding acquisition, Writing — review and editing

## Acknowledgements

We would like to thank Marius Geiss for his assistance in procuring the stimuli and in conducting some of the measurements. D.K. is supported by the Deutsche Forschungsgemeinschaft (DFG; SFB/TRR 135, project number 222641018) and an ERC Starting Grant (ERC-2022-STG 101076057). This research was further supported by “The Adaptive Mind”, funded by the Excellence Program of the Hessian Ministry of Higher Education, Science, Research and Art.

