## Supplementary Information for "Imaginary scenes are represented in cortical alpha activity"

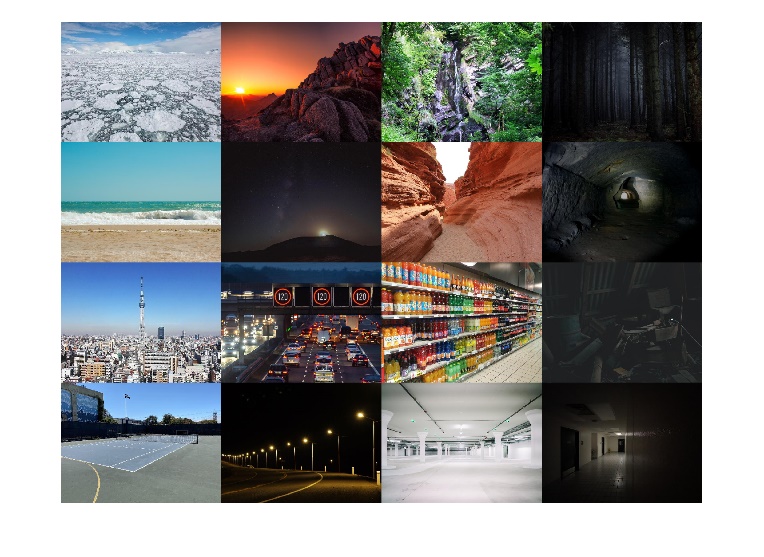

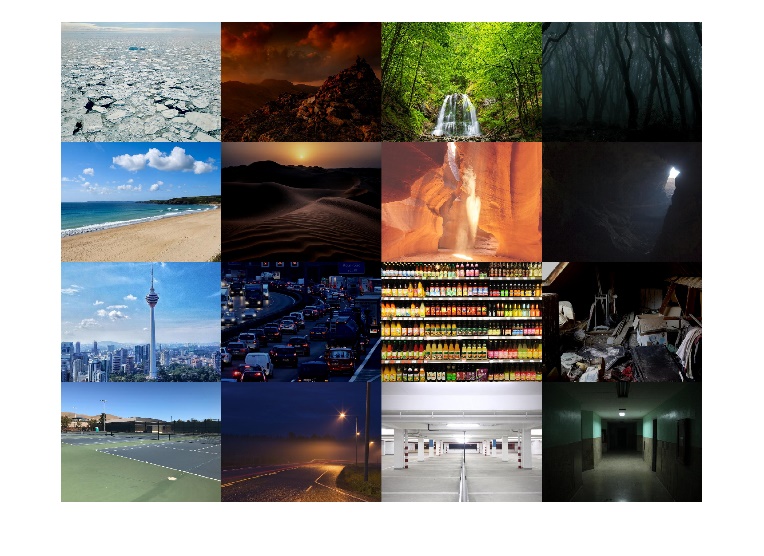

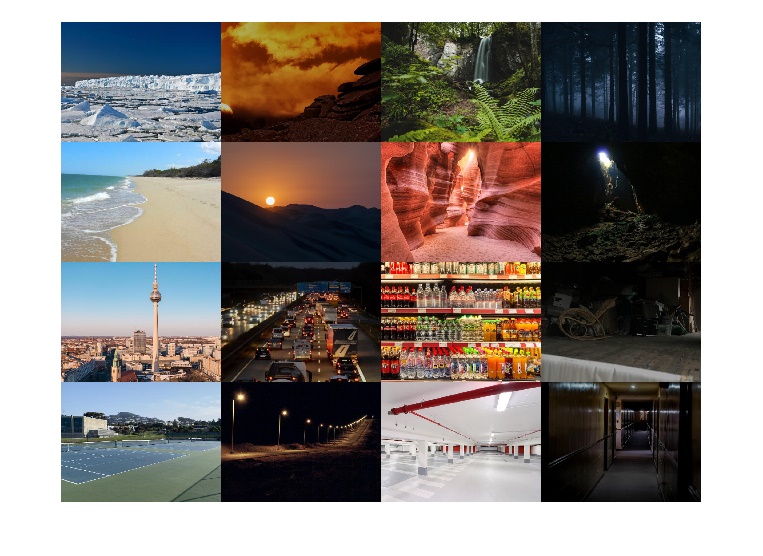

**Figure S1. Stimuli used in the perception task.**

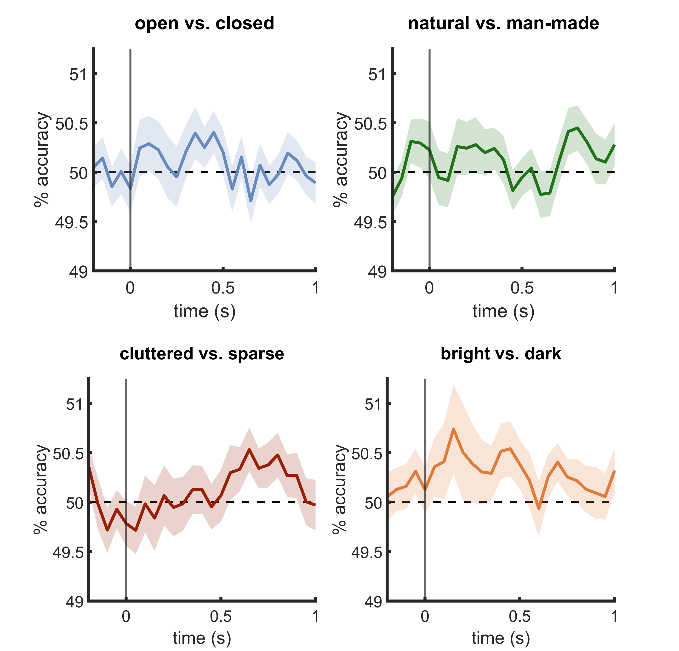

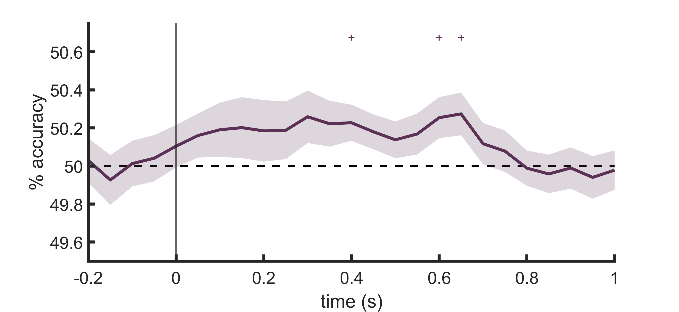

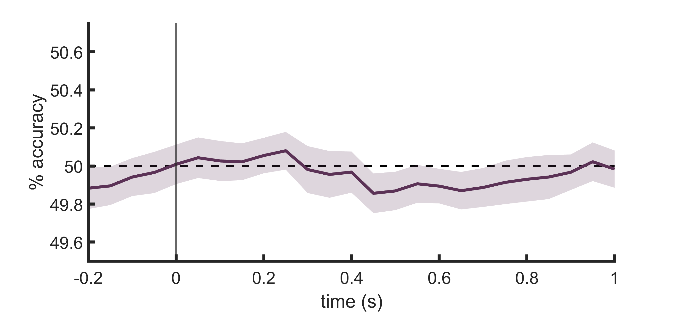

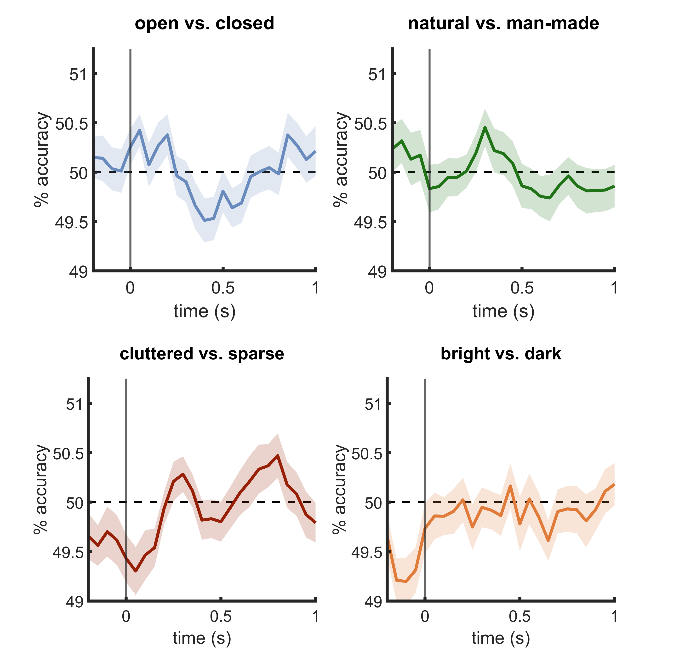

**a**

**c**

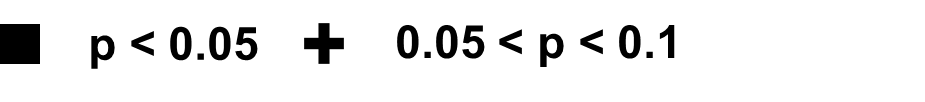

**b**

**d**

**Figure S2. Imagery-perception cross-decoding in the theta and beta frequency bands.** a) Mean pairwise scene cross-decoding in the theta band. b) Scene property cross-decoding in the theta band. c) Mean pairwise scene cross-decoding in the beta band. d) Scene property cross-decoding in the beta band. Apart from a marginally significant trend in the theta frequency band for the mean pairwise scene decoding, we found no significant above-chance cross-decoding performance. Error margins reflect the standard error of the mean. Cross markers indicate marginal significance at p<0.1 (corrected for multiple comparisons).

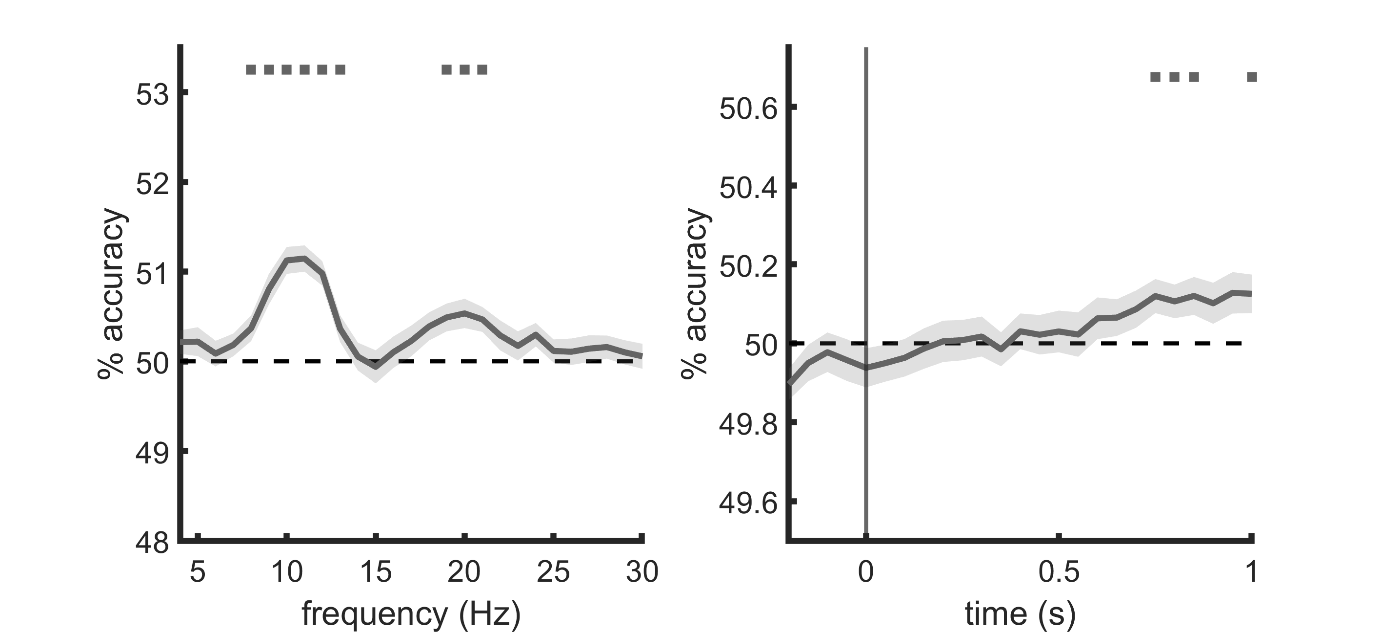

**b**

**a**

**Figure S3. Shuffled property decoding.** a) Scene property decoding with randomized property category assignment at each frequency from 4-30 Hz during imagery. b) Imagery-perception scene property cross-decoding in the alpha band with randomized property category assignment at each time point during perception. This analysis revealed late shared alpha band representations with imagery from around 750 to 850 ms and at 1000 ms during perception. These timings overlap with the marginally significant time points identified in the original mean pairwise scene cross-decoding (see Fig. 3c). Overall, both shuffled property (cross-)decoding analyses show a highly similar decoding accuracy profile with the mean pairwise scene (cross-)decoding (see Fig. 3a and 3c), suggesting that the shuffled property (cross-)decoding is indeed based on individual scene features. Error margins reflect the standard error of the mean. Square markers indicate significance at p<0.05 (corrected for multiple comparisons).

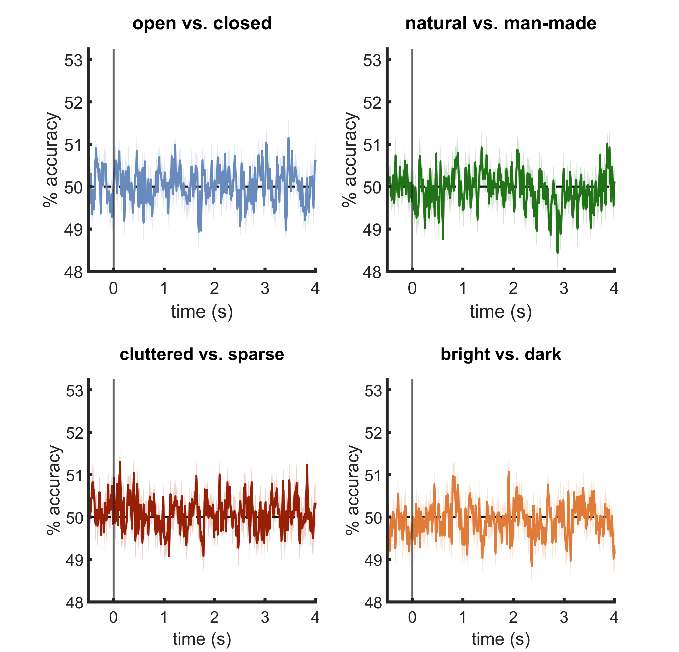

**b**

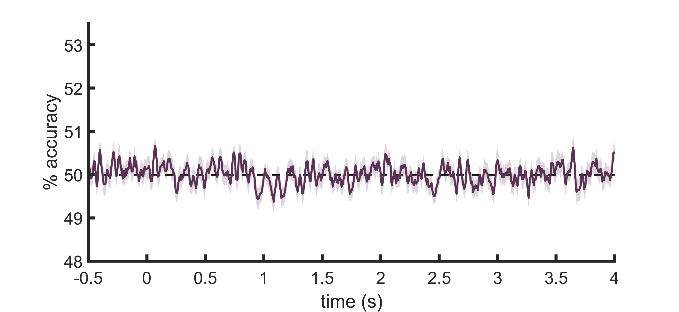

**a**

**Figure S4. Time-resolved decoding of individual imagined scenes and their properties from broadband EEG responses across all channels.** a) Mean pairwise scene decoding. b) Scene property decoding. Imagined scenes and their properties were not discriminable from broadband responses across time. Individual-participant time courses (200 Hz temporal resolution) were smoothed with a 5-time-point (25 ms) rolling average before averaging across participants. Error margins reflect the standard error of the mean.

| Openness | Naturalness | Clutter Level | Brightness | Descriptions (English) |
| --- | --- | --- | --- | --- |
| open | natural | cluttered | bright | Imagine you are on a vast arctic sea that stretches until the horizon underneath a blue sky. Your eyes survey an ocean that is filled with so many different ice floes that you can barely see the water. An infinite number of white, irregular figures, ranging from small ice chunks to large floes, are frolicking in the cold water. |
| closed | natural | cluttered | bright | Imagine you are in a deciduous forest during daytime. You are standing in front of a tall rock face from which a waterfall gushes down onto a small group of ragged rocks. You are surrounded by so many weeds, bushes and trees that you can barely see the sky, but, due to the intense daylight shining through the canopy, the forest is quite bright. |
| open | man-made | cluttered | bright | Imagine you are standing on top of an office building, looking at a gigantic urban metropolis. You are confronted with an endless sea of apartment buildings and office complexes of different sizes and shapes. A gigantic, slim TV tower lingers in the clear morning sky above the scenery. |
| closed | man-made | cluttered | bright | Imagine you are in the beverage section of a supermarket. You are looking at a shelf in the dazzling neon light that is filled with bottles from top to bottom. The bottles of different brands feature a great palette of colors and shapes and their myriad of labels indicate the beverages they contain. |
| open | natural | sparse | bright | Imagine you are at the beach. You are surprised, that the even, fine-grained sand beach is completely free of trash and that there is not a single person in sight. Your eyes rest upon a docile, blue-green sea in the glowing afternoon sun, from which every now and then small waves are being washed upon the bright sand. |
| closed | natural | sparse | bright | Imagine you are in a narrow canyon between two cliffs made of red stone. Even though the passage is so narrow that you can barely see the afternoon sky, your environment is very well lit by the sunrays. You stand on a barren, sandy ground that is completely free of rubble. |
| open | man-made | sparse | bright | Imagine you are on an empty tennis facility. The facility has blue hard courts, that are spanned by dark nets. Even though the facility is surrounded by a black fence, one can see the clear blue sky that is covered by a barren concrete building on the left very well. |
| closed | man-made | sparse | bright | Imagine you are in an underground parking house. In the dazzling light of the neon tubes on the ceiling you can see that the ceiling is painted entirely white and supported by rows of pillars. You notice in addition, that the parking house is completely empty and neither vehicles nor people are to be found. |
| open | natural | cluttered | dark | Imagine you are on a mountain ridge. A gigantic pile of bizarre jagged rocks that stretches across almost the entire ridge towers to your right. To your left, you can see the weak evening sun disappear behind the horizon, casting the scenery in a dark red. |
| closed | natural | cluttered | dark | Imagine you are in a dark forest at night. In the faint moonlight you can make out that you are fully surrounded by an impenetrable tangle of dry branches, tree trunks and loose twigs. You are also enveloped by a thick fog that makes it impossible for you to see what is ahead of you. |
| open | man-made | cluttered | dark | Imagine you are on a highway bridge in the late evening, looking down upon a traffic jam. An endless stream of densely packed cars and trucks flows slowly along the road. There is a bridge crossing the motor way right in front of you on which multiple street signs are attached. |
| closed | man-made | cluttered | dark | Imagine you are in an old and narrow attic. The attic is so dark that your environment is hard to make out. The few light beams filtering through from outside enable you to see that you are surrounded by piles of junk, consisting of old tools, furniture and boxes among other things. |
| open | natural | sparse | dark | Imagine you are in a desert at dusk. You are watching the evening sun disappear behind a giant dune made of the finest sand, dyeing the entire desert a deep black. In this almost complete darkness, you cannot make out anyone or anything around you. |
| closed | natural | sparse | dark | Imagine you are in a dark cave. In the light of your weak flashlight, you can make out that the cave is very narrow and empty and that its walls consist of blue-green stone. You also notice that there is a narrow passage behind which lies the exit of the cave, from which a tiny, distant cone of light seeps in. |
| open | man-made | sparse | dark | Imagine you are on a country road. It’s late at night and you cannot recognize anything except for the road that slithers endlessly towards the horizon and is shrouded in a soft and warm yellow. Because it is late, there are no vehicles on the road. |
| closed | man-made | sparse | dark | Imagine you are in the corridor of an office building after a power outage. The last remaining ceiling lamp casts a weak cone of light on the floor in front of you. You notice that the corridor is totally empty and there is not a single soul around you. |

**Table S1. Scene descriptions with scene property categories and English translation.** The original scene descriptions were in German and can be found on our OSF repository: https://osf.io/vxhtw/.
